# Glycine promotes longevity in *Caenorhabditis elegans* in a methionine cycle-dependent fashion

**DOI:** 10.1101/393314

**Authors:** Yasmine J. Liu, Georges E. Janssens, Rashmi Kamble, Arwen W. Gao, Aldo Jongejan, Michel van Weeghel, Alyson W. MacInnes, Riekelt H. Houtkooper

## Abstract

The deregulation of metabolism is a hallmark of aging. As such, changes in the expression of metabolic genes and the profiles of amino acid levels are features associated with aging animals. We previously reported that the levels of most amino acids decline with age in *Caenorhabditis elegans (C. elegans).* Glycine, in contrast, substantially accumulates in aging *C. elegans*. In this study we show that this is coupled to a decrease in gene expression of enzymes important for glycine catabolism. We further show that supplementation of glycine significantly prolongs *C. elegans* lifespan and ameliorates specific transcriptional changes that are associated with aging. Glycine feeds into the methionine cycle. We find that mutations in components of this cycle, methionine synthase *(metr-1)* and S-adenosylmethionine synthetase *(sams-1)*, completely abrogate glycine-induced lifespan extension. Strikingly, the beneficial effects of glycine supplementation are conserved when we supplement with serine, also driving the methionine cycle. RNA sequencing of serine- and glycine-supplemented worms reveals similar transcriptional profiles including widespread gene suppression. Taken together, these data uncover a novel role of glycine in the deceleration of aging through its function in the methionine cycle.

**Author summary:** There are a growing number of studies showing that amino acids function as signal metabolites that influence aging and health. Although contemporary -OMICs studies have uncovered various associations between metabolite levels and aging, in many cases the directionality of the relationships is unclear. In a recent metabolomics study, we found that glycine accumulates in aged *C. elegans* while other amino acids decrease. The present study shows that glycine supplementation prolongs longevity and drives a genome-wide inhibition effect on *C. elegans* gene expression. Glycine as a one-carbon donor fuels the methyl pool of one-carbon metabolism composed of folates and methionine cycle. We find that glycine-mediated longevity effect is fully dependent on methionine cycle, and that all of these observations are conserved with supplementation of the other one-carbon amino acid, serine. These results provide a novel role for glycine as a promoter of longevity and bring new insight into the role of one-carbon amino acids in the regulation of aging that may ultimately be beneficial for humans.

## Introduction

Aging is characterized by a progressive deterioration of functional capacity in tissues and organs. Pioneering studies in the nematode *C. elegans* identified longevity-associated genes and provided us with great insights into the plasticity of aging (1-3). In the last few decades, genetic and nutritional interventions have been employed in multiple organisms, including *Saccharomyces cerevisiae, C. elegans, Drosophila melanogaster*, rodents, and more recently fish (4-6). These models have set the stage for characterizing the genetic basis of physiological aging and for developing efficient strategies to control the rate of aging.

To date, metabolic pathways including the mTOR, insulin/IGF-1, and AMP-activated protein kinase (AMPK) signaling pathways emerge as being critical on aging [reviewed in (7)]. Several studies demonstrate that the levels of specific amino acids effectively influence lifespan by affecting these pathways. For example, restriction of methionine extends the lifespan of flies in a mTOR-dependent manner (8). Moreover, the branched-chain amino acids valine, leucine, and isoleucine when administered to *C. elegans* can function as signaling metabolites, that mediate a mTOR-dependent neuronal-endocrine signal that in turn promotes a longer lifespan (9). Finally, inhibition of threonine and tryptophan degradation also contributes to lifespan extension by enhancing protein homeostasis in *C. elegans* (10, 11).

In addition to the mTOR signaling pathway, alterations in one-carbon metabolism involving the folate and methionine cycle couple amino acid metabolism to the regulation of human health and disease (12). Glycine, as one of the input amino acids that feeds into one-carbon metabolism, provides a single carbon unit to the folate cycle to yield a variety of one-carbon bound tetrahydrofolates (THFs). These function as coenzymes in methylation reactions including production of methionine through methionine synthase (METR-1 in *C. elegans*) as well as the universal methyl donor, S-adenosylhomocysteine (SAMe) through S-adenosyl methionine synthetase (SAMS-1 in *C. elegans*). These output metabolites of one-carbon metabolism support a range of biological functions (13). In *C. elegans* mutations in the metabolic gene *sams-1* and the levels of SAMe and SAH have been implicated in the regulation of aging (14, 15). Although the underlying mechanism of how SAMe/SAH status influences aging needs further investigation, studies *in vivo* have provided evidence that the level of SAMe couples with the trimethylation status of lysine 4 on histone H3 (H3K4me3) and affects gene regulation (16). Another study in mouse pluripotent stem cells demonstrated that threonine catabolism contributes one carbon to SAMe synthesis and histone methylation through the glycine cleavage pathway (17). Of particular note, several histone methyl-transferases and de-methyltransferases in *C. elegans* have been identified as longevity regulators (18-20). Taken together, these studies all suggest that altering one-carbon metabolism is a mechanism to control the aging process.

We recently showed that glycine accumulates with age in a large scale metabolomics study profiling levels of fatty acids, amino acids, and phospholipids across the lifespan of *C. elegans*, (21).Another study in human fibroblast suggested that the epigenetic suppression of two nuclear-coded genes, glycine C-acetyltransferase *(GCAT)* and serine hydroxymethyltransferase 2 *(SHMT2)*, which are involved in glycine synthesis in mitochondria, was partly responsible for aging-associated mitochondrial respiration defects and that glycine treatment rejuvenated the respiration capacity of fibroblasts derived from elderly individuals (22). However, to date the role of glycine has not been systematically defined in animal models of longevity. In this study, we build upon our previous observations that suggest glycine accumulation in aging animals may play a unique and as-of-yet unexplored role in the regulation of eukaryote lifespan.

## Results

### Genes coding for glycine degradation enzymes decrease with age in *C. elegans* as glycine accumulates

We previously measured amino acid levels throughout the life of *C. elegans*, including four larval phases (L1-L4) and ten days of adulthood from young worms to aged ones (days 1-10) (21). We reported that the concentrations of most amino acids peak at the later larval stage or early adult phase and then began declining at different adult stages, reaching low levels by the latest stages of the animals’ life (21). One stark exception was glycine, which continued to accumulate in aged worms (21).

To determine if the accumulation of glycine with age is due to increased synthesis or reduced degradation, we measured the expression levels of genes directly involved in glycine metabolism in worms collected at different ages (Fig 1A). Specifically, we observed that the expression levels of most genes involved in glycine synthesis, including threonine aldolase *(R102.4)*, serine hydroxymethyltransferase *(mel-32)*, and alanine-glyoxylate aminotransferase 2(*T09B4.8*), remain unchanged in aged worms (Fig 1B). In contrast, the expression levels of genes in glycine degradation and consumption pathways, including glycine decarboxylase *(gldc-1)*, phosphoribosylamine-glycine ligase *(F38B6.4)*, and D-amino acid oxidase *(daao-1)* were dramatically lower at day 9 of adulthood (D9) (Fig 1C). This is particularly the case for *F38B6.4*, which was reduced to 6.9 % of the level at day 1 (D1) (Fig 1C). These data suggest that the accumulation of glycine observed in aged worms is predominantly due to a reduction in the expression of genes required for its degradation.

**Fig 1.**
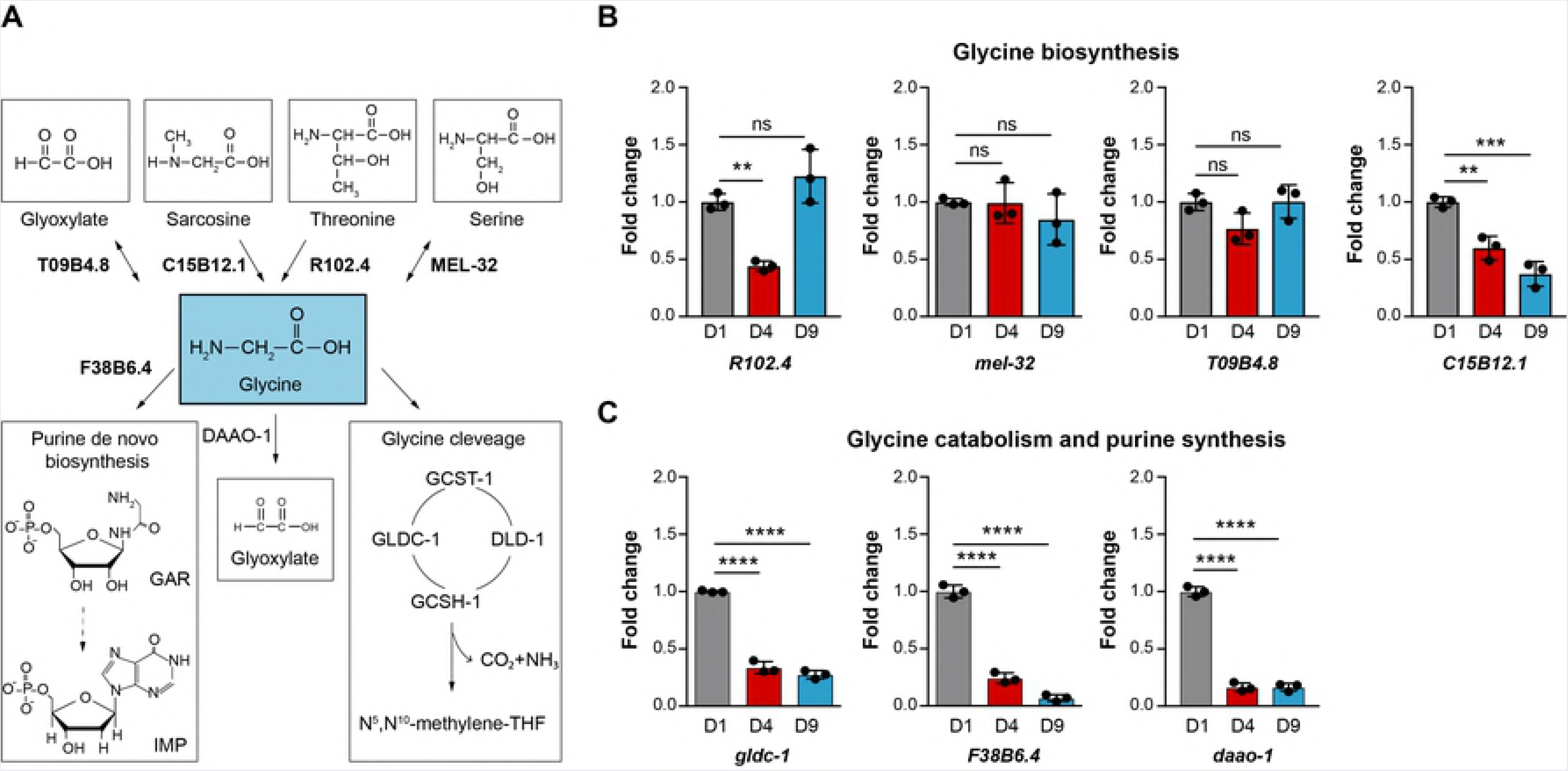
The Abundance of glycine increases with age in *C. elegans*. **(A)** Glycine metabolic pathways in *C. elegans* with genes used in RT-qPCR. This diagram was constructed according to “glycine, serine and threonine metabolism” (cel00260) in KEGG pathway database (57). **(B)** Transcript levels of genes involved in glycine biosynthesis measured by RT-qPCR showing that most genes assessed including *R102.4, mel-32*, and *T09B4.8* are quite stable throughout the life stages of worms including D1, D4, and D9. Results are shown relative to transcript levels in D1 adult worms. Mean ± SD with three biological replicates; **p < 0.01; ***p < 0.001; ns, not significant. **(C)** Transcript levels of genes involved in glycine catabolism and purine *de novo* synthesis showing that all the genes assessed including *gldc-1, F38B6.4*, and *daao-1* decrease markedly in aged worms (D9). Results are shown relative to transcript levels in D1 adult worms. Mean ± SD with three biological replicates; **p < 0.01; ***p < 0.001; ****p < 0.0001; ns, not significant.

### Dietary glycine decelerates aging in *C. elegans*

We next verified the causal involvement of glycine in lifespan regulation by determining the effects of supplementing various glycine concentrations on worm lifespan. To avoid influences of glycine on bacterial metabolism and vice versa, we effectively killed the *E. coli* OP50 with a combination of ultraviolet (UV)-irradiation and antibiotic (carbenicillin) supplementation.

In line with a previously reported observation (23), worms being fed UV-killed *E. coli* OP50 live significantly longer compared to those being fed live *E. coli* OP50 (S1 Fig). Therefore, to confirm if glycine still accumulates in aged worms after switching to this killed bacterial diet, we again quantified amino acid levels in worms at different stages of *C. elegans* lifespan including L3 (larval stage 3), day 1 (D1), day 3 (D3), day 6 (D6) and day 9 (D9) of adulthood. In this timeframe, we found that most amino acids either decreased or remained unchanged (S2A Fig), but glycine still showed a steady increase from D1 to D9 (Fig 2A). This confirmed that the accumulation of glycine is a robust phenotype that is independent of the worms being fed live or killed bacteria.

**Fig 2.**
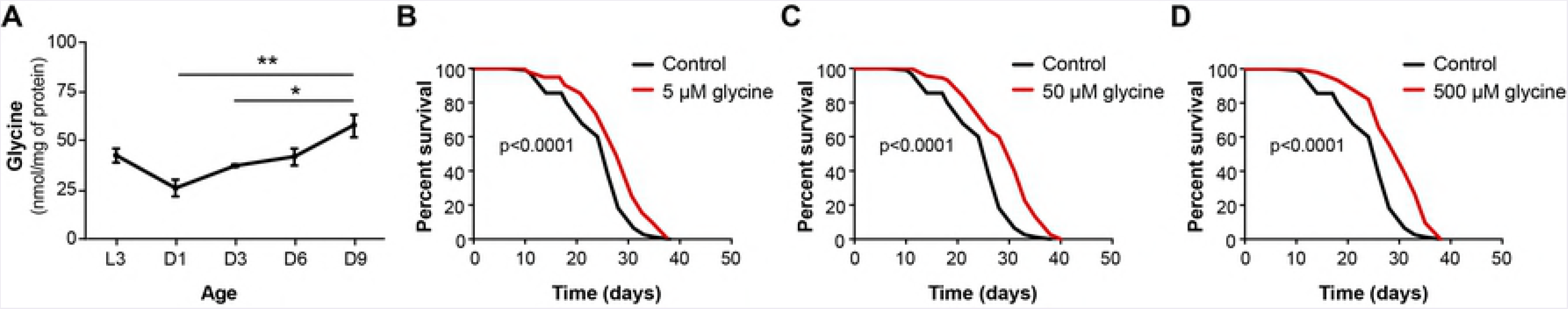
Glycine extends the lifespan of *C. elegans*. **(A)** HPLC-MS/MS quantification of glycine in life history of nematode from larval stage 3 (L3) to D9 adult worms. Levels of other amino acids measured from the same batch of samples were indicated in S2A Fig. Mean ± SD with three biological replicates; *p < 0.05; **p < 0.01. **(B-D)** Lifespan analyses of *C. elegans* cultured on UV-killed bacteria showing that glycine supplementation significantly extends lifespan at molarities of 5 μM, *p* < 0.0001 **(B)**, 50 μM, *p* * 0.0001 **(C)**, and 500 μM, *p* < 0.0001 **(D)**. Worms were exposed to dietary glycine throughout larval development and the entire adult lifespan. See S1 Table for lifespan statistics.

On UV-killed *E. coli* OP50, we tested how a range of glycine concentrations from 5 μM to 10 mM affects the lifespan of *C. elegans*. We observed a significant increase in median lifespan at concentrations of 5 μM, 50 μM, and 500 μM of dietary glycine as compared to untreated controls, with 7.7% (p < 0.0001), 19.2% (p < 0.0001), and 19.2% (p < 0.0001) extension respectively (Figs 2B-2D). Higher concentrations, however, including 5 mM and 10 mM, failed to extend worm lifespan suggesting a dose response where only low concentrations of glycine between 5-500 μM are beneficial to lifespan (Figs 2B-2D, S2B and S2C Figs). Our findings are in agreement with previous studies suggesting a dose-effect of amino acids on worm lifespan (24).

Next, we measured amino acid levels in *C. elegans* at D1 of the adult stage to investigate how glycine supplementation at different doses affected glycine levels *in vivo*. We did not detect obvious changes of glycine abundance itself or any of the other amino acids at any concentration of supplementations (S3A-S3D Figs). These results suggest that in worms, glycine immediately fuels metabolic pathways and that the levels of amino acids are tightly controlled in the early adult stage.

### Glycine supplementation counteracts aging-induced repression of one-carbon and purine metabolism Genes

We next sought to understand how one-carbon and purine metabolism are dynamically influenced by aging. We therefore quantified the expression levels of genes involved in one-carbon metabolism (Fig 3A) at 5 time points (L3, D1, D3, D6 and D9) throughout the worm lifespan. We found that with the exception of *tyms-1* (thymidylate synthetase) and *dhfr-1* (dihydrofolate reductase), which are upregulated in aged worms, genes participating in transferring the one-carbon moiety of glycine to form SAMe, including *gcst-1* (glycine cleavage system T-protein), *dld-1* (dihydrolipoamide dehydrogenase), *gcsh-1* (glycine cleavage system H-protein), *mthf-1* (methylene tetrahydrofolate reductase), *metr-1* (methionine synthase), *and sams-1* (S-adenosyl methionine synthetase), dropped dramatically in aged animals (Fig 3B). Furthermore, the purine *de novo* synthesis genes *atic-1* (5-aminoimidazole-4-carboxamide ribonucleotide formyltransferase/IMP cyclohydrolase) (Fig 3B) and *F38B6.4* (Fig 1C) were markedly reduced in aged worms compared to their expression levels in worms at D1. Together these data suggest a downregulation of both one-carbon and purine metabolism during aging in *C. elegans*.

**Fig 3.**
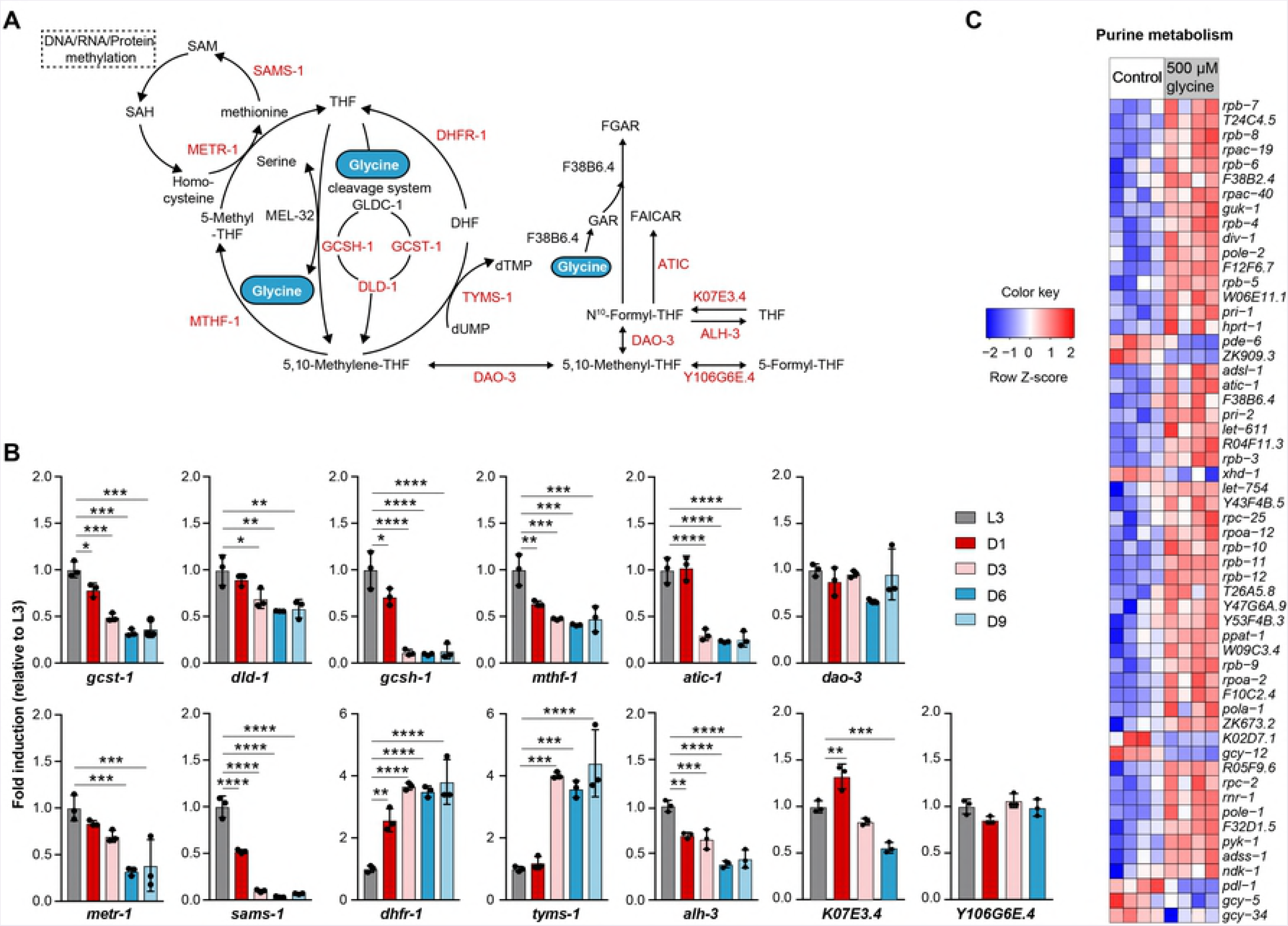
Glycine supplementation counteracts the age-induced suppression of gene expression in purine and one-carbon metabolism. **(A)** Metabolic network of glycine metabolism feeding into one-carbon metabolism and purine *de novo* synthesis. Genes quantified by RT-qPCR and presented in the heat maps are highlighted in red. **(B)** The relative expression levels of the glycine metabolizing genes in worms from larval stage 3 (L3) to D9 adult. The expression level of the indicated gene was measured by RT-qPCR on total mRNA isolated from synchronized worms at every time point and compared to the mean in L3 worms. Mean ± SD. with three biological replicates; *p < 0.5; **p < 0.01; ***p < 0.001; ****p < 0.0001; ns, not significant. **(C)** Heat map showing transcriptional profile of the differentially expressed genes in purine metabolism upon supplementation with 500 μM glycine (z-score normalized). An adjusted *p*-value < 0.05 was set as the cut-off for significantly differential expression.

To resolve in a greater detail how glycine supplementation counteracts the age-related changes in one-carbon and purine metabolic genes, we performed next generation sequencing on UV-killed *E. coli* OP50-fed worms (control) and 500 μM glycine supplemented worms at the first day of adulthood (Fig 3C and S4 Fig). In contrast to the changes occurring with age, glycine induced a marked increase in the expression of genes in purine metabolic pathways (Fig 3C). Concomitantly several genes in one-carbon metabolism (indicated in red in Fig 3A) were differentially expressed, including upregulation of *atic-1, F38B6.4, dhfr-1, tyms-1, mel-32*, and *gcsh-1*, and downregulation of *sams-1, mthf-1*, and *gldc-1* (S4 Fig). These results suggest that exogenous supplementation of dietary glycine compensates for the loss of one-carbon and purine synthesis metabolic activity, directly impeding this aspect of the aging process.

### Transcriptional upregulation of glycine and one-carbon metabolism is a shared signature of longevity

Having determined the effect of glycine on worm lifespan and its ability to counteract age-related decline occurring in the one-carbon and purine metabolic pathways, we next aimed to investigate whether similar gene regulatory events were also present in long-lived mutant worms. To test this, we turned to microarray datasets from three long-lived worm models, *daf-2(e1370)* and *eat-2(ad456)* (25) or *mrps-5* RNAi worms, reported here. We specifically look at glycine-associated metabolic pathways in the Kyoto Encyclopedia of Genes and Genomes (KEGG) database including ‘ glycine serine and threonine metabolism’, ‘ one carbon pool by folate’, ‘ cysteine and methionine metabolism’, and ‘ purine metabolism’. Strikingly, although distinct longevity pathways are known to be active in these long-lived worms (2, 26, 27), all these longevity worm models consistently showed a transcriptional activation of glycine metabolism, folate-dependent one-carbon metabolism and methionine metabolism (Figs 4A-4C). This suggests that regulation of glycine and one-carbon metabolism is a life-extending mechanism. In contrast, the overall purine metabolism, including *de novo* synthesis, the salvage pathways, and purine degradation was mildly deactivated across the three long-lived strains (Figs 4A-4C).

**Fig 4.**
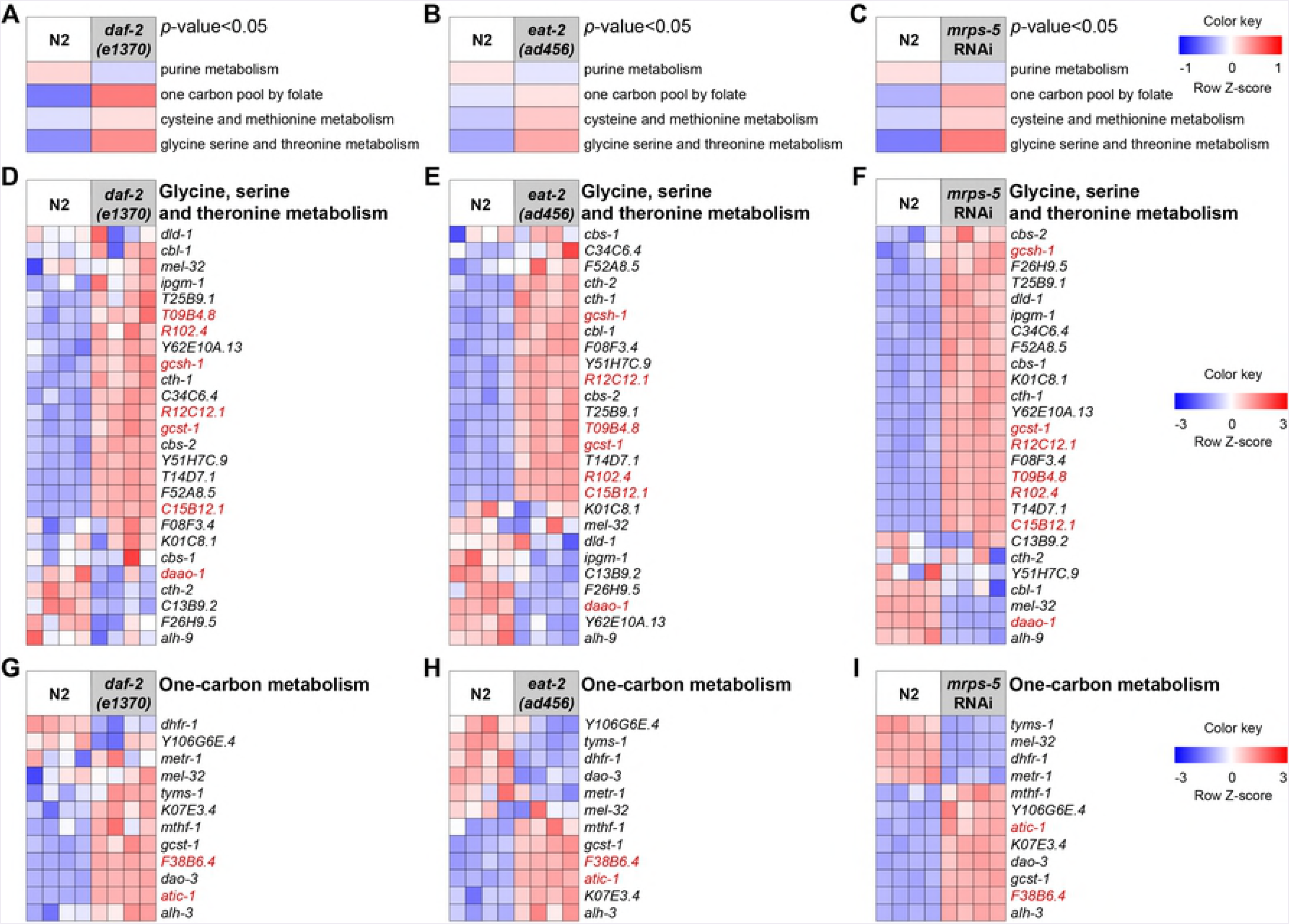
Transcriptomic changes of glycine and one-carbon metabolism in *daf-2(e1370), eat-2(ad456)*, and *mrps-5* RNAi worms. (A-C) Gene set map analyses performed on the KEGG gene sets including “glycine, serine and threonine metabolism” (cel00260), “cysteine and methionine metabolism” (cel100270), “one carbon pool by folate” (cel100670), and “purine metabolism” (cel100230) showing that these four KEGG gene sets are significantly upregulated with a p-value < 0.05 in *daf-2 (e1370)* **(A)**, *eat-2 (ad456)* **(B)** and *mrps-5* RNAi worms **(C)** compared to wild type N2 respectively. Gene set map analyses were performed on “R2” platform and plotted in heat maps (z-score normalized). A p-value < 0.05 was used as the cut-off for differentially affected gene sets in every pair of comparison. **(D-F)** Heat maps showing that the majority of genes in “glycine, serine and threonine metabolism” from the KEGG gene sets are upregulated in *daf-2 (e1370)* (D), *eat-2 (ad456)* **(E)** and *mrps-5* RNAi worms **(F)** compared to N2 respectively (z-score normalized). **(G-I)** Heat maps showing that the majority of genes in “one carbon pool by folate” from the KEGG gene sets are upregulated in *daf-2 (e1370)* **(G)**, *eat-2 (ad456)* **(H)** and *mrps-5* RNAi worms **(I)**.

To identify prominent transcriptional features in the three long-lived strains, we next examined the expression profile of individual genes belonging to “glycine, serine and threonine metabolism”, as well as folate-mediated one-carbon metabolism and purine metabolism. We found that most genes involved in glycine anabolism including *T09B4.8, C15B12.1* and *R102.4* were upregulated in *daf-2(e1370), eat-2(ad456)* and *mrps-5* RNAi worms (Figs 4D-4F). Moreover, a concomitant rise in the levels of genes in glycine catabolism and purine *de novo* synthesis occurred, including *R12C12.1, gcst-1, gcsh-1, F38B6.4*, and *atic-1* (Figs 4D-4I), suggesting a stimulation in metabolic activity of glycine-associated processes in these long-lived worms. Interestingly, the expression level of *daao-1*, which encodes the protein that unidirectionally catabolizes glycine to glyoxylate (Fig 1A), decreased in all long-lived worms, whereas the level of *T09B4.8*, which codes for the protein that produces glycine from glyoxylate, increased (Figs 4D-4F). This indicates that glycine was predominantly consumed by one-carbon metabolism and purine *de novo* synthesis. In contrast to an overall upregulation of purine metabolism genes in response to dietary glycine treatment, all three longevity worm models specifically induced the expression of genes in purine *de novo* synthesis compared to control, including *F38B6.4, F10F2.2 (pfas-1*, phosphoribosylformylglycinamidine synthase,*), B0286.3 (pacs-1*, phosphoribosylaminoimidazole succinocarboxamide synthetase), and *atic-1* (S5A-S5C Figs). Taken together, our data reveal that transcriptional activations of glycine and glycine-associated pathways, including one-carbon and purine *de novo* synthesis, are present in three distinct longevity models.

### Dietary glycine causes large-scale changes in gene expression

To understand the mechanism of glycine-mediated lifespan extension on a more global scale, we returned to our next-generation sequencing dataset and performed unsupervised Principle Component Analysis (PCA) on the individual RNA-seq libraries. We found clear separation between glycine-treated versus untreated samples (Fig 5A), corresponding to a large difference in gene expression (Fig 5B). Surprisingly, comparing genome-wide expression pattern in control and glycine-treated worms we noted a striking trend that more genes were transcriptionally repressed in response to glycine treatment (Fig 5B). Quantification of this observation confirmed that the proportion of differentially downregulated genes was 2.7-fold more than the number of differentially upregulated genes (72.79% versus 27.21%), suggesting an inhibition propensity of glycine on gene expression (Fig 5C).

**Fig 5.**
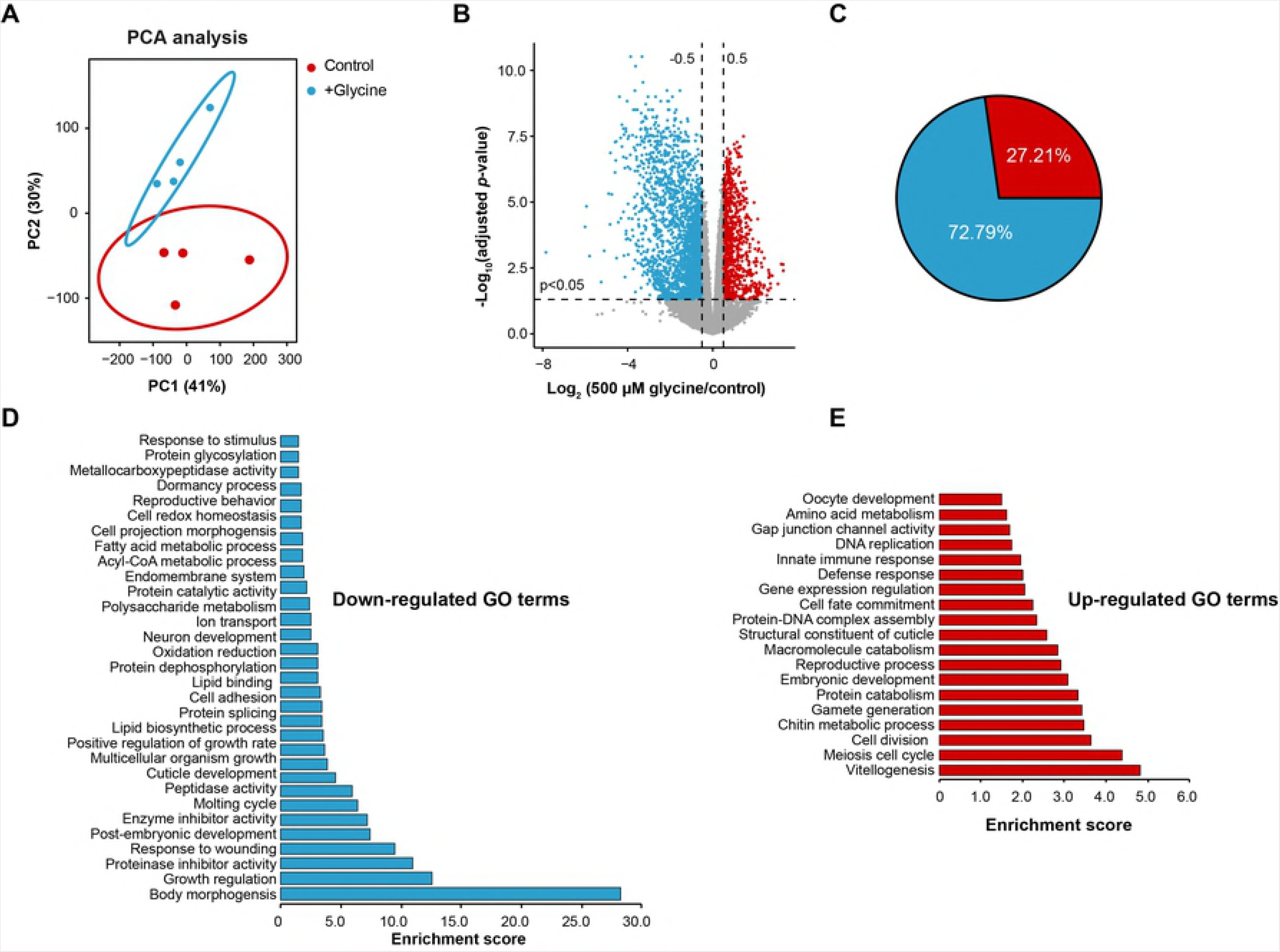
Genome-wide analyses identify cellular metabolic processes affected by glycine supplementation. **(A)** Principle Component Analysis (PCA) showing group separation based on differentially expressed genes in 500 μM glycine supplemented worms compared to control. **(B)** Differentially expressed genes in 500 μM glycine over control as showed in the volcano plot. An adjusted p-value < 0.05 and |log_2_ (500 μM glycine / control ratio) | > 0.5 were applied to determine significantly up and downregulated genes (red and blue dots), or genes without change in expression (grey dots). **(C)** The percentages of genes with a significant increase or decrease in expression in the total number of differentially expressed genes showed respectively in the pie chart. An adjusted *p*-value < 0.05 and |log_2_ (500 μM glycine / control ratio) | > 0.5 were used to determine differentially expressed genes. Red part indicates the proportion of upregulated genes, blue indicates the proportion of downregulated genes. **(D and E)** Gene ontology (GO) term enrichment analyses (biological processes) of the 2629 significantly downregulated genes **(D)** and the 973 significantly upregulated genes **(E)** (blue and red dots from the volcano plots B with a p-value < 0.05) performed using DAVID Bioinformatics Database with an EASE score < 0.05.

To probe the processes changed upon glycine supplementation, we performed gene ontology (GO) term enrichment analysis on the differentially expressed genes using the Database for Annotation, Visualization and Integrated Discovery (DAVID) bioinformatics resource (28). A larger number of GO terms were found enriched in the downregulated gene list, among which were GO terms for body morphogenesis, growth regulation, post-embryonic development, molting cycle, cuticle development, multicellular organism growth, and positive regulation of growth rate (Fig 5D). These enrichments all related to growth control, suggesting a decelerating effect of glycine upon development and growth which is a phenomenon known to be associated to longevity (29). Additionally, unlike some mutations conferring longevity to the soma at the cost of a reduction in fecundity (30), supplementation of glycine mildly enhanced the expression of genes in reproduction-related biological processes, including GO terms of vitellogenesis, meiosis cell cycle, and gamete generation (Fig 5E). Overall, with the observations of the lifespan extending effects of glycine, these observations suggest that the beneficial role of glycine slows down pathways that are traditionally ameliorated in healthy aging models.

### Methionine cycle integrity is required for glycine-mediated lifespan extension

By fueling the one-carbon metabolic network glycine contributes to the synthesis of SAMe, which is the major methyl carrier in all methylation events of nucleic acids, lipids, and proteins such as histones (Fig 3A). This led us to hypothesize that the methionine cycle may be required for the lifespan-extending effect of glycine. To test if the methionine cycle is necessary for the longevity effect of glycine, we performed lifespan analyses with the methionine cycle-deficient mutants *metr-1(ok521)* and *sams-1(ok3033)*. In these mutants, 500 μM glycine failed to promote lifespan (Figs 6A-6C), demonstrating that the effects of glycine on worm longevity depend on the methionine cycle.

**Fig 6.**
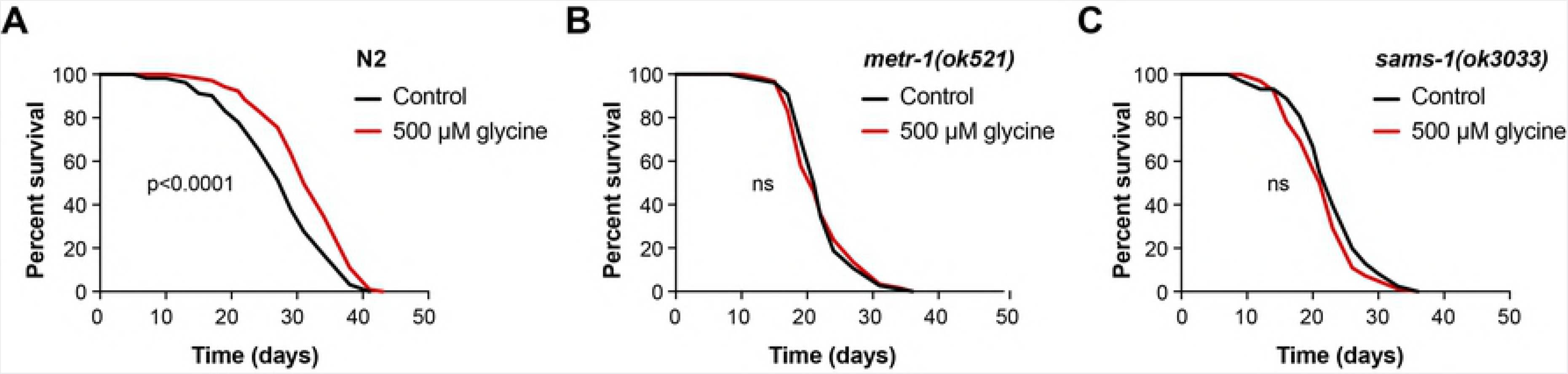
Methionine cycle is required for the longevity effect of glycine in *C. elegans*. **(A)** The lifespan analysis of N2 worms fed control diet (UV-killed OP50) or 500 μM glycine diet showing that 500 μM glycine supplementation significantly promotes lifespan extension (p < 0.0001).**(B and C)** Lifespan analysis of the mutant worms of *metr-1(ok521)* animals **(B)** or *sams-1(ok3033)* animals **(C)** fed 500 μM glycine diet compared to their corresponding mutants fed control diet (UV-killed OP50). See S1 Table for lifespan statistics.

### Serine shares the conserved mechanism with glycine to extend lifespan

Serine is another important one-carbon donor and the major precursor for glycine synthesis *in vivo* (13). Hydroxymethyltransferase (MEL-32 in *C. elegans*) converts serine into a one-carbon unit to form *N^5^-* N^10^-methylene tetrahydrofolate and glycine that can be further catabolized by the glycine cleavage system (Fig 3A). Thus, serine and glycine are closely related to each other in one-carbon metabolism. Given their similarities, we next investigated whether serine can also exert beneficial effects on worm lifespan. Serine supplementation at a concentration from 1 mM to 10 mM has been shown previously to extend worm lifespan (24). We therefore administered 5 mM serine to worms and measured lifespan. We confirmed that serine prolongs the lifespan of worms (+20.8% in median lifespan) (Fig 7A). We further investigated whether the anti-aging effects of serine also relied on the methionine cycle. In agreement with our results observed with glycine supplementation, the lifespan extending effect of serine was ablated by mutations of *metr-1* and *sams-1* as well (Figs 7B and 7C), suggesting serine and glycine prolong *C. elegans* lifespan via a similar mechanism.

**Fig 7.**
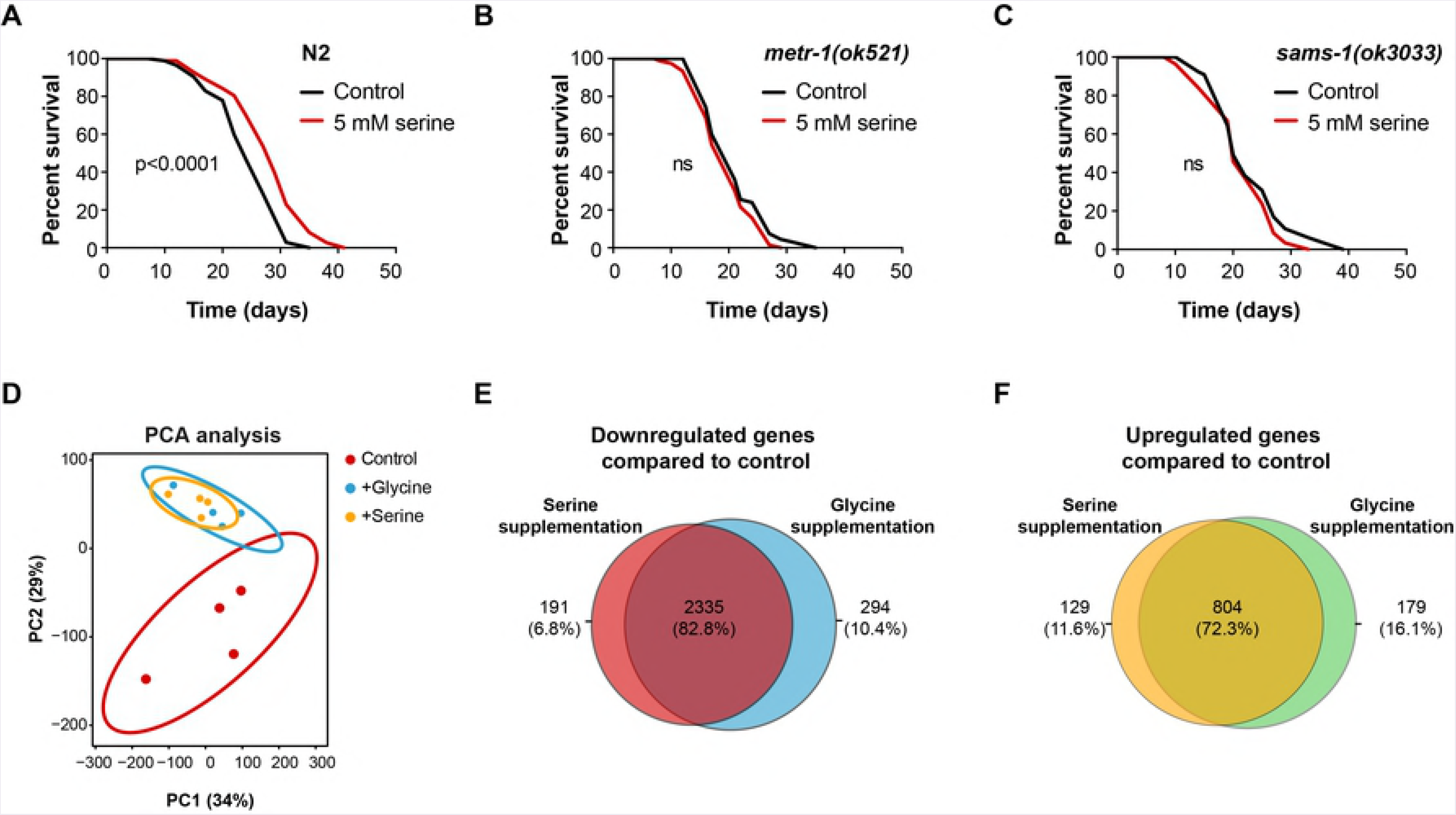
Serine increases *C. elegans* lifespan in a methionine cycle-dependent fashion. **(A)** Lifespan analysis of N2 showing that 5 mM serine promotes longevity (p < 0.0001). **(B and C)** Lifespan analysis performed in *metr-1(ok521)* **(B)** and *sams-1(ok3033)* mutant animals **(C**) fed 500 μM glycine diet compared to their corresponding mutants fed control diet (UV-killed OP50). See S1 Table for lifespan statistics. **(D)** Principle Component Analysis (PCA) plot showing group separation based on transcriptional profiles in worms supplemented with 500 μM glycine and 5 mM serine compared to control. **(E and F)** Venn diagram comparing overlap between genes significantly downregulated or upregulated in glycine and serine supplementation groups compared to control group. 2335 downregulated genes **(E)** and 804 upregulated genes **(F)** were found to be shared by glycine and serine supplementation groups. An adjusted p-value < 0.05, |log_2_ (500 μM glycine / control ratio)| > 0.5, |log_2_ (5 mM serine / control ratio)| > 0.5 were used to determine differentially expressed genes.Supporting Information Lege

To determine the common regulators in both glycine and serine-mediated longevity, we performed genome-wide RNA-seq on serine-supplemented worms. As expected, PCA analysis showed clear separation between worms treated with either of the amino acids when compared to non-treated worms, and a strong similarity between glycine- and serine-treated worm groups (Fig 7D). Statistical analysis found one significantly differentially expressed gene, *F38B6.4* (S6A Fig), an enzyme that consumes glycine for purine synthesis. In addition, visualizing the data as a volcano plot showed again a greater number of genes repressed by serine treatment, similar to what was observed with glycine (S6B Fig). The number of genes downregulated in worms supplemented with serine compared to controls was found to be roughly 2.7 times greater than the number of upregulated genes, which follows the same gene expression suppression pattern of glycine supplemented worms (S6C Fig). Likewise, we found a strong overlap between glycine- and serine-treated worms when looking at the up- and downregulated genes, as shown in the Venn diagrams where 82.8% (2335) of the downregulated and 72.3% (804) of the upregulated genes are shared (Figs 7E and 7F). Taken together, these results support the conclusions that both glycine and serine supplementation feed into the methionine cycle to induce suppression of gene expression and stimulate longevity through common signaling pathways.

## Discussion

Our work sheds light onto the means by which the amino acid glycine can increase *C. elegans* lifespan when supplemented to the diet. Using a metabolomics approach, we found that glycine steadily and significantly accumulates in aging *C. elegans*. Furthermore, we demonstrated that this accumulation is mainly coupled to a decrease in the expression levels of genes in glycine cleavage and purine synthesis pathways. We found that dietary glycine reverses these transcriptional changes and extends lifespan at concentrations between 5-500 μM, while mutations in methionine synthase *[metr-1(ok521)]* and S-adenosyl methionine synthetase *[sams-1 (ok3033)]*, two enzymes involved in methionine cycle, can fully abrogate this lifespan extension. Moreover, we found that serine, another amino acid that feeds into one-carbon metabolism, shows similar transcriptional changes, *metr-1* and *sams-1* dependency, and lifespan extension upon dietary supplementation as does glycine. These results confirm an important role for the methionine cycle in the longevity effects of glycine.

Our work identified a counterintuitive biological phenomenon, whereby glycine accumulation was observed during the aging process in worms while supplementation of glycine was nonetheless able to prolong worm lifespan. However, it is not uncommon for changes that occur with age to also benefit lifespan when artificially induced. For example, suppression of IGF1 signaling may extend lifespan in many model organisms (2, 31, 32), while IGF1 levels themselves have been observed to decline with age (33, 34). Moreover, methionine restriction is beneficial to lifespan in a variety of model organisms (8, 35), while methionine abundance *in vivo* has been observed to decline with age [observed in this study (S2 Fig) and (21)]. Similar to these phenomena, glycine supplementation may activate protective cellular pathways that promote longevity when exogenously applied, while a natural glycine accumulation with age may reflect the organism’s need to upregulate these same cytoprotective pathways to deal with the damage and detrimental changes occurring during aging.

We have shown a conserved upregulation of genes in glycine and one-carbon metabolism in long-lived worms, including *daf-2(e1370), eat-2(ad456)*, and *mrps-5* RNAi. This observation suggests that intervention in glycine-related metabolic processes is the common feature in diverse longevity programs and can be activated by simple glycine supplementation. While many core transcriptional features associated with longevity have been described in recent decades, including the downregulation of insulin/IGF signaling, suppression of mTOR signaling, and upregulation of the mitochondrial unfolded protein response (UPR^mt^), our work suggests glycine-related one-carbon metabolism to be a novel core transcriptional feature through which genetic and nutritional intervention prevent aging. These observations warrant further investigation, to see if classical longevity pathways are dependent fully or in part on glycine metabolism for their lifespan benefiting effects.

Studies in rodents have suggested glycine supplementation to be pro-longevity (36, 37), to have anti-inflammatory effects (38), to be cytoprotective (39), and to ameliorate metabolic disorders (40). In humans, glycine supplementation in patients with metabolic disorders has a protective effect against oxidative stress and inflammation (41-43). In line with these observations in mammalian systems, our data demonstrated that glycine promotes longevity in *C. elegans*. Furthermore, we show this benefit to occur in a *metr-1* and *sams-1* dependent manner, implicating the methionine cycle in longevity regulation. Finally, we show glycine supplementation induces widespread suppression of genes including many that are hallmarks of the aging process. Taken together, our findings suggest that dietary glycine is an effective strategy to increase lifespan and warrant further investigation for life- and health-span studies in humans.

## Materials and methods

### Worm strains and maintenance

*C. elegans* strains N2 Bristol, RB2204 *sams-1(ok3033)X* and RB755 *metr-1(ok521)II* were obtained from the Caenorhabditis Genetics Center (CGC, University of Minnesota). Nematodes were grown and maintained on Nematode growth media (NGM) agar plates at 20°C as previously described (44).

### Bacterial feeding strains and RNAi experiments

*E. coli* OP50 was obtained from the CGC, *mrps-5* RNAi bacterial clone was derived from the Vidal library (45). Bacterial feeding RNAi experiments were carried out as described (46). Synchronized eggs were placed onto plates seeded with HT115 and *mrps-5* RNAi bacteria respectively. At the young adult stage, worm samples were collected for mRNA extraction.

### Preparation of amino acid supplementation plates

Glycine and serine were purchased from Merck Millipore (no. 8.1603.0250) and Sigma (no. S4500), respectively. A stock of concentration of 1 M glycine and serine was made by dissolving glycine and serine in water. The pH of glycine and serine stock solution was adjusted to 6.0-6.5 with sodium hydroxide and then sterilized with 0.45 μm Millipore filter. The concentrations of 5 μM, 50 μM, 500 μM, 5 mM and 10 mM glycine, and 5 mM serine were used in the present study.

### Preparation of UV-killed bacteria

Overnight cultures of *E. coli* OP50 were seeded on standard NGM plates containing carbenicillin (25 μg ml^-1^) to prevent the bacterial growth. After drying overnight at room temperature, the bacterial lawn was irradiated with 254 nm UV light using a Stratalinker UV Crosslinker model 1800 (Stratagene, USA) at 999900 μJ/cm^2^ for 5 min. A sample of UV-exposed *E. coli* OP50 was collected and cultured in LB medium overnight at 37°C to confirm the bacteria were completely killed. Plates seeded with UV-killed bacteria were stored in 4 °C and used within 1 week after seeding.

### Lifespan measurements

Lifespan experiments were performed at 20°C without fluorouracil, as described (47). Briefly, amino acid supplementation started at egg stage and 120-150 worms per condition were used for lifespan. During the reproductive period (≈ day 1-8), worms were transferred to fresh plates every day to separate them from their progeny. Survival was scored every other day throughout the lifespan and a worm was considered as dead when they did not respond to three taps. Worms that were missing, displaying internal egg hatching, losing vulva integrity and burrowing into NGM agar were censored. Statistical analyses of lifespan were calculated by Log-rank (Mantel-Cox) tests on the Kaplan-Meier curves in GraphPad Prism.

### Amino acid extraction and UPLC-MS/MS analysis

Amino acids were extracted and analyzed as described before (21) and each experiment was performed in three biological replicates. Briefly, around 1500 synchronized worms at the desired stage were harvested, freeze-dried and stored at room temperature until use. Worm lysates were obtained by homogenization and subsequent tip sonication. Amino acids were extracted from worm lysate containing 50 μg protein and measure by UPLC-MS/MS analysis.

### Extraction of mRNA and quantitative real-time PCR (qPCR) for gene expression in *C. elegans*

Approximately 500 worms were collected in three biological replicates per condition. Total RNA was isolated according to the manufacturer’ s protocol. Briefly, samples were homogenized in TRIzol (Invitrogen) with a 5 mm steel metal bead and shaken using a TissueLyser II (Qiagen) for 5 min at a frequency of 30 times/sec. RNA was quantified with a NanoDrop 2000 spectrophotometer (Thermo Scientific) and stored at −80 °C until use.Genomic DNA was eliminated, and cDNA was synthesized using the QuantiTect Reverse Transcription kit (QIAGEN). The qPCR reaction was carried out in 8 μL with a primer concentration of 1 μM and SYBR Green Master mix (Roche) in a Roche LightCycler 480 system. In all analyses, the geometric mean of two reference genes, *eif-3.C* and F35G12.2, was used for normalization and the oligonucleotides used for PCR are listed in S2 Table.

### Microarray

Microarray experiment was performed as described (25). Approximately 500 worms were collected in four replicates per condition and total RNA was extracted as described above. RNA quality and quantity were assessed after DNase clean-up using a 2100 Bioanalyzer (Agilent Technologies). RNA was amplified and labeled using a Low Input QuickAmp Labeling Kit (Agilent Technologies) and hybridized using the Agilent Gene Expression Hybridization Kit (Agilent Technologies). An ArrayXS-068300 with WormBase WS241 genome build (OakLabs) was used and fluorescence signals were detected by the SureScan microarray Scanner (Agilent Technologies). Data of all samples were quantile normalized using the ranked median quantiles as described previously (48).

### Samples for RNA library preparation

Approximately 500 worms were collected in quadruplicates per condition for total RNA extraction as described above. Genomic DNA residues were eliminated with RNase-Free DNase (Qiagen), followed with the cleaning up with the RNeasy MinElute Cleanup Kit (Qiagen). Samples were sent to GenomeScan B.V. (Leiden, The Netherlands) for RNA library preparation and sequencing at a 20 million read-depth (see methods below).

### RNA library preparation and next-generation sequencing (RNA-seq)

Samples were processed for Illumina using the NEBNext Ultra Directional RNA Library Prep Kit (NEB #E7420) according to manufacturer’s description. Briefly, rRNA was depleted using the rRNA depletion kit (NEB# E6310). A cDNA synthesis was performed in order to ligate with the sequencing adapters. Quality and yield after sample preparation was measured with the Fragment Analyzer. Size of the resulting products was consistent with the expected size distribution (a broad peak between 300-500 bp). Clustering and DNA sequencing using the Illumina cBot and HiSeq 4000 was performed according to manufacturer’s protocol with a concentration of 3.0 nM of DNA. HiSeq control software HCS v3.4.0, image analysis, base calling, and quality check was performed with the Illumina data analysis pipeline RTA v2.7.7 asnd Bcl2fastq v2.17.

### RNA-seq: read mapping, statistical analyses, and data visualization

Reads were subjected to quality control FastQC (http://www.bioinformatics.babraham.ac.uk/projects/fastqc), trimmed using Trimmomatic v0.32 (49) and aligned to the *C. elegans* genome obtained from Ensembl, wbcel235.v91 using HISAT2 v2.0.4 (50). Counts were obtained using HTSeq (v0.6.1, default parameters) (51) using the corresponding GTF taking into account the directions of the reads. Statistical analyses were performed using the edgeR (52) and limma/voom (53) R packages. All genes with no counts in any of the samples were removed whilst genes with more than 2 reads in at least 4 of the samples were kept. Count data were transformed to log2-counts per million (logCPM), normalized by applying the trimmed mean of M-values method (Robinson et al., 2010) and precision weighted using voom (54). Differential expression was assessed using an empirical Bayes moderated t-test within limma’s linear model framework including the precision weights estimated by voom (54). Resulting *p*-values were corrected for multiple testing using the Benjamini-Hochberg false discovery rate. Genes were re-annotated using biomaRt using the Ensembl genome databases (v91). RNA-seq samples were compared using principal component analysis (PCA) and Partial least squares discriminant analysis (PLS-DA) using mixomics (55). Heatmaps, venn diagrams, and volcano plots were generated using ggplot2 (56) in combination with ggrepel (https://CRAN.R-project.org/package=ggrepel), and venneuler (https://CRAN.R-project.org). Data processing and visualization was performed using R v3.4.3 and Bioconductor v3.6.

### Functional annotation

Gene ontology (GO) analyses were conducted using DAVID bioinformatics resource (28). Genes found to be significantly up- or downregulated with an adjusted p-value < 0.05 and | log_2_ (fold change) | > 0.5 were subjected to functional annotation clustering. To retrieve significantly enriched GO terms, enrichment threshold (EASE score) was set as 0.05 for all analyses and the category of each annotation cluster generated by David was curated manually.

### Gene expression profile visualization

Heat maps of gene expression profile in Fig 4 and S5 Fig were plotted using “R2”, Genomics analysis and Visualization platform (http://r2.amc.nl). Gene set map analyses were performed on “R2” with defined gene sets from the Kyoto Encyclopedia of Genes and Genomes (KEGG) pathways (http://www.genome.jp/kegg/) (57).

### Quantification and statistical analysis

Data were analyzed by two-tailed unpaired Student’s t-test or by one-way ANOVA with Tukey’s post hoc test for multiple comparisons, except for survival curves, which were calculated using the log-rank (Mantel-Cox) method. For all experiments, data are shown as mean ± SD and a p-value < 0.05 was considered significant.

### Data availability

The microarray data is available at Gene Expression Omnibus (accession number: GSE117581). The RNA-seq datasets discussed in this publication are available at the NCBI Sequence Read Archive (SRA) (accession number: SRP155700).

## Acknowledgments

The authors thank the Caenorhabditis Genetics Center at the University of Minnesota, which is funded by NIH Office of Research Infrastructure Programs (P40 OD010440), for providing *C. elegans* strains. Work in the Houtkooper group is financially supported by an ERC Starting grant (no. 638290), and a VIDI grant from ZonMw (no. 91715305). AWM is supported by E-Rare-2, the ERA-Net for Research on Rare Diseases (ZonMW #40-44000-98-1008). GEJ is supported by a Federation of European Biochemical Society (FEBS) long-term fellowship.

## Author contributions

YJL and RHH conceived and designed the project. YJL, RK and AWG performed experiments. GEJ and AJ performed bioinformatics. YJL, GEJ, AWM, and RHH interpreted data. YJL, GEJ, AWM and RHH wrote the manuscript, with contributions from all other authors.

## Conflict of interest

The authors declare that they have no conflict of interest.

---

**S1 Fig.Culturing worms on UV-killed bacteria extends the lifespan of *C. elegans*** Lifespan analysis of *C. elegans* cultured on alive and UV-killed *E. coli* OP50, showing that the latter live longer. See S1 Table for lifespan statistics.

**S1 Fig. Culturing worms on UV-killed bacteria extends the lifespan of *C. elegans.***

Lifespan analysis of *C. elegans* cultured on alive and UV-killed *E. coli* OP50, showing that the latter live longer. See S1 Table for lifespan statistics.

**S2 Fig. Amino acids profiles change with age in *C. elegans.***

**(A)** Amino acid amounts were measured using UPLC-MS/MS across 5 time-points of the lifespan of worms fed UV-killed bacteria, including L3, D1, D3, D6, and D9. Note: the profile of glycine was shown in Fig 2A. Statistical analysis between groups was performed using one-way ANOVA, followed by Tukey’s post hoc test for multi-comparisons and significance levels were indicated with asterisks as follows: *p < 0.05; **p < 0.01, ***p < 0.001. **(B and C)** Lifespan analysis of *C. elegans* cultured on UV-killed bacteria upon 5 mM **(B)** and 10 mM **(C)** glycine treatment.

**S3 Fig. Exogenous glycine supplementation didn’t alter amino acid profile in day 1 adult worms.**

**(A-D)** Amino acid profile of D1 adult worms supplemented with increasing amounts of glycine ranging from 5 μM to 10 mM and fed UV-killed OP50. The levels of glycine, serine, and methionine in **(A)** were presented in separate graphs **(B)**, **(C)**, and **(D)** with the levels of significance indicated. Statistical analysis was performed using one-way ANOVA, followed by multi-comparisons with Tukey’s post hoc test to compare the mean of glycine treated groups with the mean of control respectively. Significance levels were indicated with asterisks as follows: ns, not significant; *p < 0.05; **p < 0.01.

**S4 Fig. Glycine supplementation counteracts the age-induced suppression of gene expression in one-carbon metabolism**

Heat map showing transcriptional changes of the genes in the Fig 3A diagram upon supplementation with 500 μM glycine (z-score normalized). Genes that are differentially expressed in worms supplemented with 500 μM glycine are highlighted in red. An adjusted *p*-value < 0.05 was set as the cut-off value for significantly differential expression.

**S5 Fig. Gene expression profile in purine metabolism in long-lived worms.**

**(A-C)** Genes in KEGG “purine metabolic pathway” in *C. elegans* were all included in the analysis. Previously reported microarray data of long-lived worms^39^ was used to compare *daf-* 2 *(e1370)* **(A)**, *eat-2(ad456)* **(B)** and *mrps-5* RNAi worms **(C)** to N2 respectively for transcriptional changes, visualized in heat map (expression values were normalized by z-score transformation) (GEO accession number: GSE106672).

**S6 Fig. Genome-wide transcriptome comparison reveals that more genes are deactivated in *C. elegans* in response to serine treatment.**

**(A)** Differentially expressed genes in 5 mM serine treated worms against 500 μM glycine treated worms as showed in Volcano plot. Grey dots indicate no differential regulation, red and blue dots indicate significant activation and repression based on adjusted p-values < 0.05 and log_2_-transformed fold change with absolute value > 0.5.

**(B)** Differentially expressed genes in 5 mM serine compared to control displayed in volcano plot. Grey dots indicate no differential regulation, red and blue dots indicate significant activation and repression based on adjusted p-values < 0.05 and log_2_-transformed fold change with absolute value > 0.5.

**(C)** Quantification for the percentage of activated and repressed genes against the total number of differentially expressed genes, represented as red and blue dots in the volcano plot showing that 73.03% was downregulated while 26.97% was upregulated, plotted in pie chart.

**S1 Table**. Summary of mean lifespan and statistical analysis (*p*-values) for lifespan experiments including different treatments and bacterial conditions displayed in **Figs 2B-2D, 6A-6C, 7A-7C, S1, S2B** and **S2C Figs**. Larval stage 4 (L4) is considered as day 0 of the lifespan assay. The mean lifespan and p-values were calculated by a log-rank (Mantel-Cox) statistical test comparing glycine or serine treatment population to the water control. *P*-values less than 0.05 are considered statistically significant, demonstrating that the two lifespan populations are different. Cumulative statistics and statistics of individual experiments are shown for each condition. The total number of individuals scored, and independent
experiments are shown.

**S2 Table. List of primers used for qPCR in *C. elegans***

